# Comparison of intracellular and encapsidated viral RNA reveals selective constraints on reovirus recombination

**DOI:** 10.64898/2026.10.10.757791

**Authors:** Alejandra Flores, Andrew Routh, Kristen M. Ogden

**Affiliations:** Department of Pathology, Microbiology, and Immunology, Vanderbilt University Medical Center, Medical Center North, 1161 21st Ave. South, Suite D-2220, Nashville, TN 37232, USA; ClickSeq Technologies LLC, 630 Pena Drive, Suite 100, Davis, CA 95618, USA; Department of Pediatrics, Vanderbilt University Medical Center, 2200 Children’s Way, Suite 2404, Nashville, TN 37232, USA

## Abstract

Recombination can generate genetic diversity among RNA viruses; yet, mechanisms governing recombination for segmented, double-stranded RNA viruses remain poorly understood. Reovirus genomes contain ten segments of double-stranded RNA. Previous analyses suggested low rates of reovirus RNA recombination and led to a proposed mechanism of recombination during minus-strand synthesis. These analyses focused on recombination events detected in genomic double-stranded RNA extracted from purified viral particles. In the current study, we looked deeper into the origins of the observed recombination events by assaying total cellular material, which comprises viral genomic material, as well as replication intermediates, mRNAs, and products of recombination that may not be eligible for packaging into virus particles. Intracellular viral RNA, which is almost exclusively plus-strand RNA, exhibited substantially higher recombination frequency than particle-associated double-stranded RNA. The diversity and frequency of identified junctions were more similar among samples of a given type (intracellular or virion-encapsidated) from distinct infections than between distinct sample types from the same infection. Although potential recombination sites are widely distributed across the genome, a more limited set was used in junctions in packaged viral RNA than in intracellular viral RNA. Together, these observations suggest recombination may occur more frequently than previously recognized during reovirus infection, and many such events likely occur during plus-strand synthesis. However, recombination products generated during viral transcription may be selectively incorporated into viral particles.

**Summary:** Recombination is an important means for RNA viruses to adapt and evolve, but recombination mechanisms for segmented, double-stranded RNA viruses are poorly understood. In previous work, we investigated recombination events in packaged viral RNA for the double-stranded RNA virus reovirus. We detected recombination relatively infrequently, and we proposed a mechanism of recombination during genomic RNA synthesis. Here, we report that recombination events are more frequent and potential recombination sites are used less selectively in viral plus-strand RNA transcripts isolated from infected cells than in viral double-stranded RNA packaged in matched purified virus particles. These findings suggest recombination is more promiscuous and widespread for double-stranded RNA viruses than previously recognized. However, the tightly coordinated assortment and packaging mechanisms of these viruses may selectively filter recombination products in the viral population.

## Introduction

Recombination generates genetic diversity in RNA viruses, contributing to evolution, host adaptation, and immune escape [1-5]. While extensively characterized for positive- and negative-strand RNA viruses, recombination mechanisms remain poorly understood for segmented double-stranded RNA (dsRNA) viruses [4, 6, 7]. Reoviruses contain ten dsRNA segments: three large (L), three medium (M), and four small (S) [8]. Within host cells, transcription and genome replication occur inside reovirus core particles. During transcription, the minus strand of each segment is the template for synthesis of plus-strand RNAs, which are released into the cytoplasm for protein synthesis or packaged into assembling viral cores. There, they become templates for complementary minus strand synthesis to generate the dsRNA genome. Previous studies identified low reovirus recombination frequencies and showed that junctions preferentially occur at short regions of microhomology [9, 10]. Although microhomology sequences are widespread throughout the viral genome, only a small subset are detected in recombination sites in packaged viral RNA [10]. These observations support a model in which recombination is highly selective. However, these studies analyzed RNA from purified viral particles and, therefore, characterized only the subset of recombined RNAs that persisted through assembly and packaging.

Like many segmented RNA viruses, reoviruses employ selective packaging mechanisms [11-18]. Evidence suggests reovirus packages one of each of the 10 gene segments or none [17]. Thus, gene segment assortment and packaging are highly ordered. However, in some cases, reovirus packages a defective gene segment [9, 10, 19-21]. Defective viral genomes (DVGs) are truncated versions of the genome generated by recombination; some DVGs retain terminal cis-acting elements required for replication and packaging, allowing incorporation into viral particles [10, 22-26]. DVGs lacking cis-acting elements are inefficiently packaged [23, 25-28]. Consequently, recombination products recovered from purified virions may represent a selectively retained subset rather than the complete population generated during infection.

In the current study, we assayed recombination events in total cellular nucleic acids, including encapsidated genomic viral dsRNA, as well as replication intermediates, mRNAs, and recombination products that may not be eligible for packaging into virus particles. We found that intracellular viral RNA contains a distinct and more diverse population of recombination products than particle-encapsidated RNA, supporting a model in which recombination products are generated under moderate constraints during plus-strand RNA synthesis and subjected to additional selection, likely during assortment and packaging.

## Results

### Characterization of reovirus recombination junction frequency

To compare the diversity of recombination junctions generated during reovirus infection, we conducted ClickSeq-based high-throughput RNA sequencing. We included three virus clones in two independent experiments. Each experiment comprised particles (Particles 1.1–3.1 and 1.2–3.2) and matched infected cells (Infected Cells 1.1–3.1 and 1.2–3.2), maintaining direct sample pairing. Infections were limited to approximately one round of replication (24 h) but conducted at a high multiplicity of infection (10 PFU/cell). At the endpoint, a portion of cells was harvested for immediate RNA extraction, with the remainder used for gradient purification of reovirus particles. Purified particles were treated with benzonase to remove non-encapsidated nucleic acids before RNA extraction. RNA was used to generate ClickSeq libraries, which were sequenced on an Illumina NovaSeq platform [29].

We obtained high-quality data for all 10 segments but focused analyses on two large segments (L1 and L3) and two medium segments (M1 and M3) to provide a snapshot of the reovirus genome. Sequence depth was consistently maintained within four-log_10-_ to six-log_10-_fold coverage across samples, segments, and replicates **(S1 Fig)**. Observed coverage biases likely reflect variability in priming, reverse transcription, amplification efficiency, and intrinsic RNA sequence features. Coverage was sufficient for downstream analyses.

Recombination junctions were identified using a viral recombination mapping algorithm (ViReMa) and defined as insertions or deletions greater than 5 bp, flanked on both sides by at least 25 bp of high-quality alignment [10, 30, 31]. Recombination junctions were detected across all segments, with junction counts generally corresponding to read counts (**S1 Table**). Intracellular viral RNA consisted almost exclusively of plus-strand RNA (98.5 ± 0.5%), while particle-associated RNA contained recombination junctions from both RNA strands, with minus-strand junctions slightly predominating (55.2 ± 7.3%) (**S1 Table**). Particle-associated RNA exhibited a mean junction frequency (JFreq; recombination-containing reads per 10,000 mapped reads) of 9.3 (SD = 4.0) **(S1 Table)**. Infected-cell samples displayed a substantially higher average JFreq of 46.1 (SD = 7.0) (**S1 Table**). Thus, recombination junctions are present at higher frequencies in intracellular than particle-encapsidated viral RNA.

### Quantitative comparison of recombination junction profiles between particle and infected cell samples

To compare the identities and distributions of recombination junctions between sample populations, we used principal coordinate analysis (PCoA), which is useful for complex data with many zeros and rare variables. We generated a Bray-Curtis dissimilarity matrix to capture differences in junctional identity and relative abundance and used PCoA to project these relationships into a reduced-dimensional space (**Fig. 1**). Across all analyzed segments, particle-associated and intracellular viral RNA typically separated along the first principal coordinate (PCoA1), indicating recombination profiles differed more between sample types than matched samples from the same infection (**Fig. 1**). Infected-cell samples consistently formed relatively tight clusters, whereas particle samples exhibited greater dispersion, particularly along the second principal coordinate (PCoA2). Despite some segment-specific differences, the patterns indicate that recombination profiles are more similar within sample type than between distinct sample types derived from the same infection.

**Figure 1.**
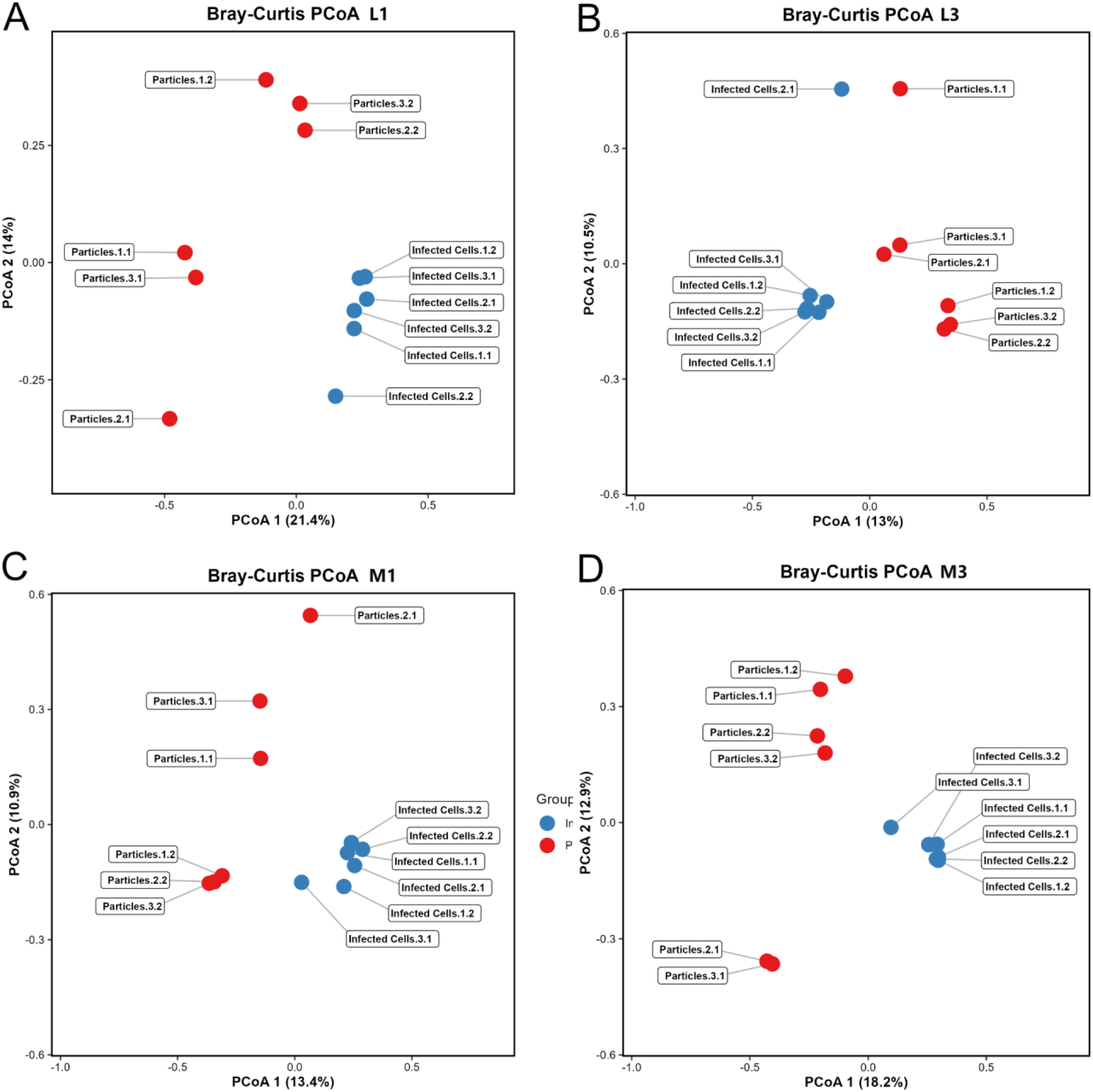
Principal coordinate analysis of reovirus recombination profiles. Score plots for principal coordinate analyses (PCoA) of gene segments L1 (A), M1 (B), L3 (C), and M3 (D) applied to junctional recombination frequency data. Data represent reovirus particles and infected cells from two independent experiments, each with three biological replicates per condition.

### Microhomology patterns differ between particle-associated and intracellular viral RNA

To compare microhomology patterns between RNA from particles and infected cells, we extracted and positionally aligned 50 nucleotides surrounding recombination junction start and stop sites for segments L1, L3, M1, and M3, quantified nucleotide identity, and normalized it to total junctions. With four possible nucleotides, by chance identity should be ∼25%. In particle samples, high positional identity was observed within ∼9 nucleotides upstream and ∼3 nucleotides downstream of the junction (**Figs. 2A and S2A**). Strand-specific analyses in particle-encapsidated RNA suggest variable microhomology around plus-strand junctions, with the strongest identity from -3 to +2 (**S3A Fig**), whereas minus-strand junctions exhibited higher identity from -9 to +3 (**S3B Fig**). For infected-cell samples, positional identity was largely confined to the −3 to -1 positions, with lower overall identity (**Figs. 2A and S2A**). Consistent with these trends, junctions more frequently contained microhomology regions in particle samples. For segments L1, L3, M1, and M3, ∼17-33% of junctions in particle samples contained six identical nucleotides, and ∼28-47% contained four identical nucleotides surrounding recombination junctions (**Tables 1 and S2**). Infected-cell samples showed markedly lower six-nucleotide (∼1-5%) and four-nucleotide (∼6-11%) junctional identity. These results suggest viral RNA in particles tends to contain recombination junctions at microhomology sites, while junctional microhomology is less prevalent in intracellular viral RNAs.

**Figure 2.**
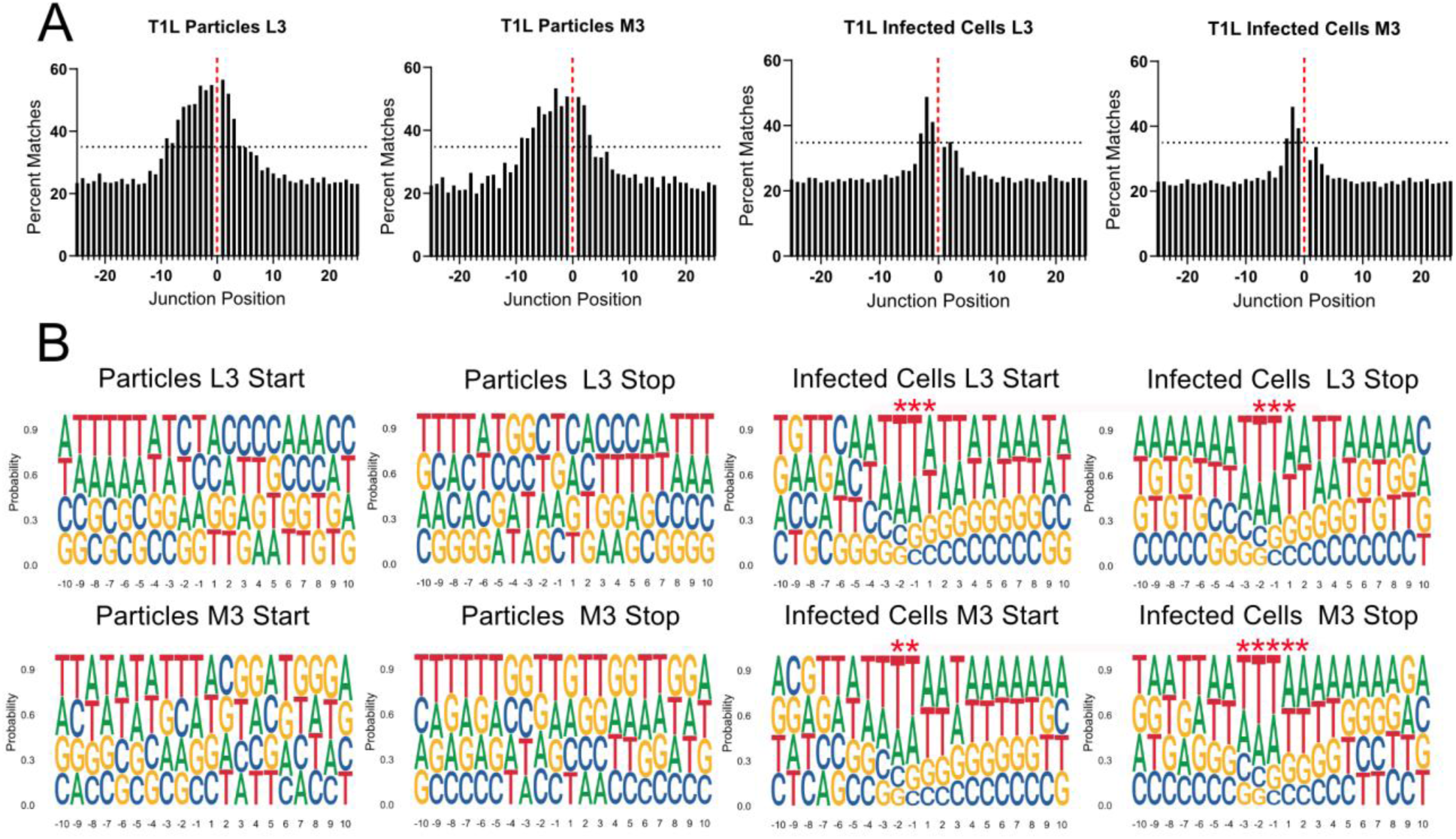
Microhomology and logoplots. (A) Sequence microhomology surrounding recombination junctions in particle and infected-cell viral RNA. The nucleotides that overlap at each position in 50-bp regions flanking the start and stop recombination junction sites were quantified for segments L3 and M3 and are expressed as a percentage of the total positions at any detected recombination junction. The junction is labeled zero and denoted with a dashed red line. The black dashed line denotes y = 35. (B) Logoplots showing nucleotides surrounding recombination junctions in particle and infected-cell viral RNA. Nucleotide composition was calculated as the percent adenosine (A), cytosine (C), guanine (G), and thymine (T) at each position in a 20-bp region surrounding the start and stop recombination junction sites. Positions upstream (−10 to −1) and downstream (+1 to +10) of the junction position are indicated. Red asterisks denote statistical significance via a Chi-Square Goodness-of-Fit test against an expected uniform baseline distribution and a threshold at which the dominant nucleotide constitutes at least 35% of the total local population. Each analyzed condition contains combined data from two independent experiments, each comprising three biological replicates. Sequence logo and microhomology analyses were conducted using all unique recombination junctions detected in each condition: L3 particles (n = 1,936), L3 infected cells (n = 4,460), M3 particles (n = 655), and M3 infected cells (n = 4,147).

To compare nucleotide preferences between RNA from particles and infected cells, we analyzed nucleotide frequencies for recombination junctions in segments L1, L3, M1, and M3. Consistent with previous observations [9, 10], combined and strand-specific analyses of particle-associated RNA showed no nucleotide enrichment at positions flanking the junctions (**Figs. 2B, S2B, and S3C-D**). Infected-cell samples displayed a modest but consistent nucleotide bias immediately surrounding the junction and corresponding with the microhomology regions (**Figs. 2 and S2**). Across all segments, thymine (uridine in RNA) was enriched at −3 to −1, followed by weaker enrichment of adenine. Adenine was enriched at +1, followed by weaker enrichment of thymine (uridine in RNA). While recombination in particle-associated RNA appears sequence-independent, intracellular viral RNA exhibits a slight nucleotide bias near junction sites, suggesting sequence context may contribute modestly to recombination.

### Microhomology sites are differentially utilized during recombination

To determine whether intracellular conditions influence recombination site selection, we identified all unique four-nucleotide microhomology sequences at recombination junction start and stop sites in segments L1, L3, M1, and M3 (**Figs. 3 and S4**). We quantified total occurrences of each sequence within the corresponding segments (**Figs. 3A and S4A**). Consistent with previous observations [10], four-nucleotide sequences were broadly distributed across segments, typically occurring ∼10-20 times. We assessed how frequently each sequence was used in recombination events in particle or infected-cell RNA samples (**Figs. 3B and S4B**). In particle samples, most four-nucleotide sequences (78–89%) were not used in any detected junction, some (10-18%) contributed to a single recombination event, and a small fraction (1–5%) were used multiple times.

**Figure. 3.**
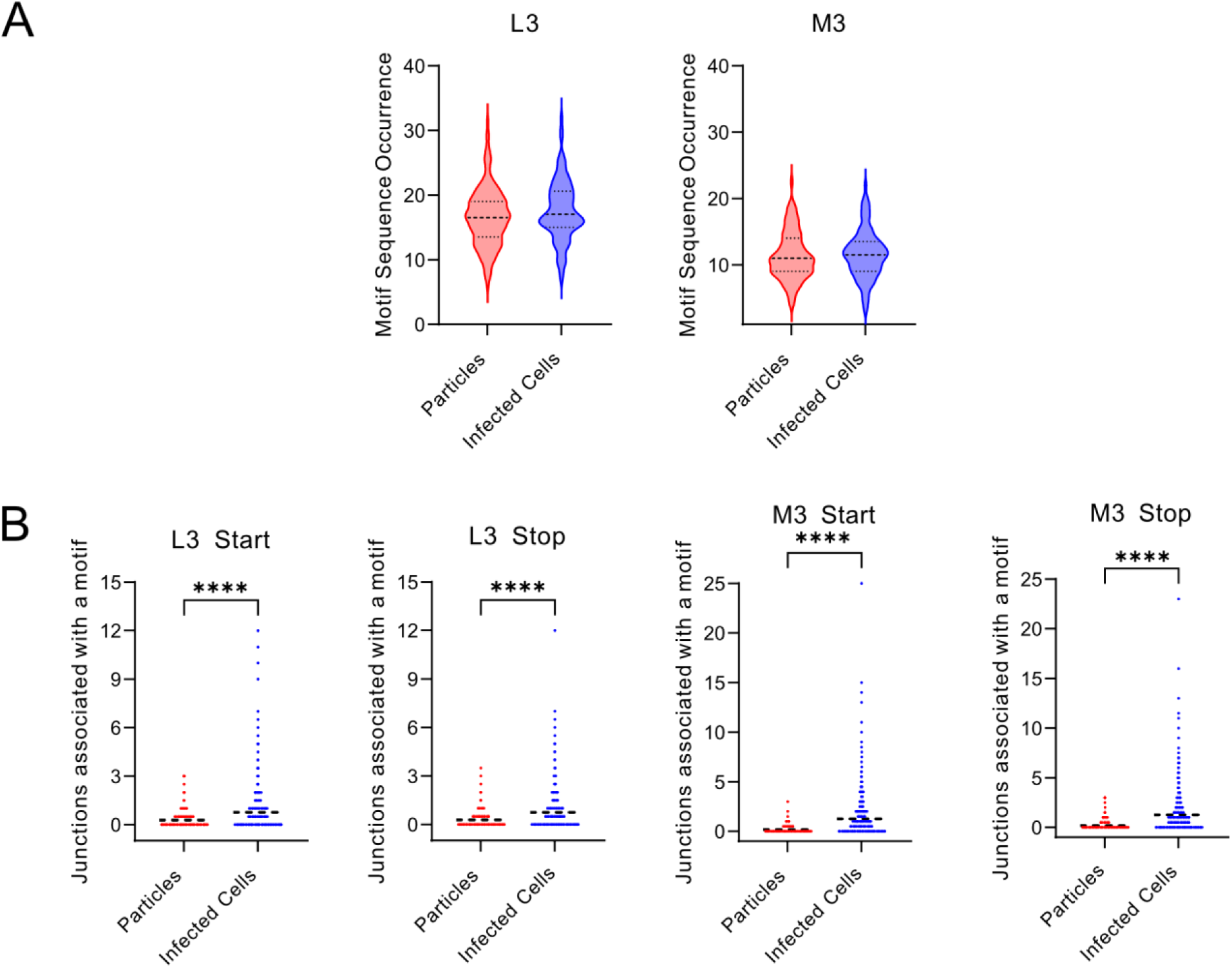
Recombination site selection. (A) Frequency of microhomology sequences in segments. The number of times each unique four-nucleotide microhomology sequence identified at a recombination junction is present in segment L3 and M3 was computationally quantified. For particles, L3: n = 337 and M3: n = 163 unique four-nucleotide microhomology sequences. For infected cells, L3: n = 181 and M3: n = 162 unique four-nucleotide microhomology sequences. Dashed lines represent the median and quartiles. (B) Usage of microhomology sites for recombination. The number of times each occurrence of a four-nucleotide sequence from a unique start site (B) or stop site (C) analyzed in (A) was incorporated into a recombination site in the L3 or M3 segment computationally quantified for particles and infected cells. Each analyzed condition contains combined data from two independent experiments, each comprising three biological replicates. For particles, L3: n= 5664 and M3: n= 1858 instances. For infected cells, L3: n= 3223 and M3: n= 1895 instances. Dashed lines represent the mean.

Infected-cell samples showed significantly broader utilization of available microhomology sequences, with fewer remaining unused (53–75%), a higher proportion contributing to a single recombination event (17–23%), and an increased frequency of repeated usage (8-24%). Thus, based on analysis of similar numbers of viral reads, intracellular viral RNA more frequently utilizes available microhomology sequences than particle-encapsidated RNA.

## Discussion

We compared recombination events in encapsidated genomic viral dsRNA with replication intermediates, mRNAs, and recombination products that may not be eligible for packaging into virus particles. Profiles of recombination junction location and frequency were more similar among samples of a given type (intracellular or encapsidated) than between distinct sample types from the same infection (**Fig. 1**). The increased recombination frequency and utilization of microhomology sequences in largely plus-strand intracellular viral RNA compared with encapsidated viral dsRNA suggest recombination occurs during viral transcription and is less selective than previously thought; the Jfreq for intracellular reovirus RNA is on the order of MERS coronavirus, a positive-sense single-stranded RNA virus (**S1 Table and Fig. 3**) [9, 10, 30]. Sample type-specific and strand-specific differences in microhomology surrounding junctions and a modest nucleotide preference in intracellular plus-strand RNA hint at potential mechanistic differences in recombination during distinct viral replication steps (**Fig. 2 and Table 1**). These observations underscore our incomplete understanding of recombination mechanisms for dsRNA viruses like reovirus.

**Table 1:**
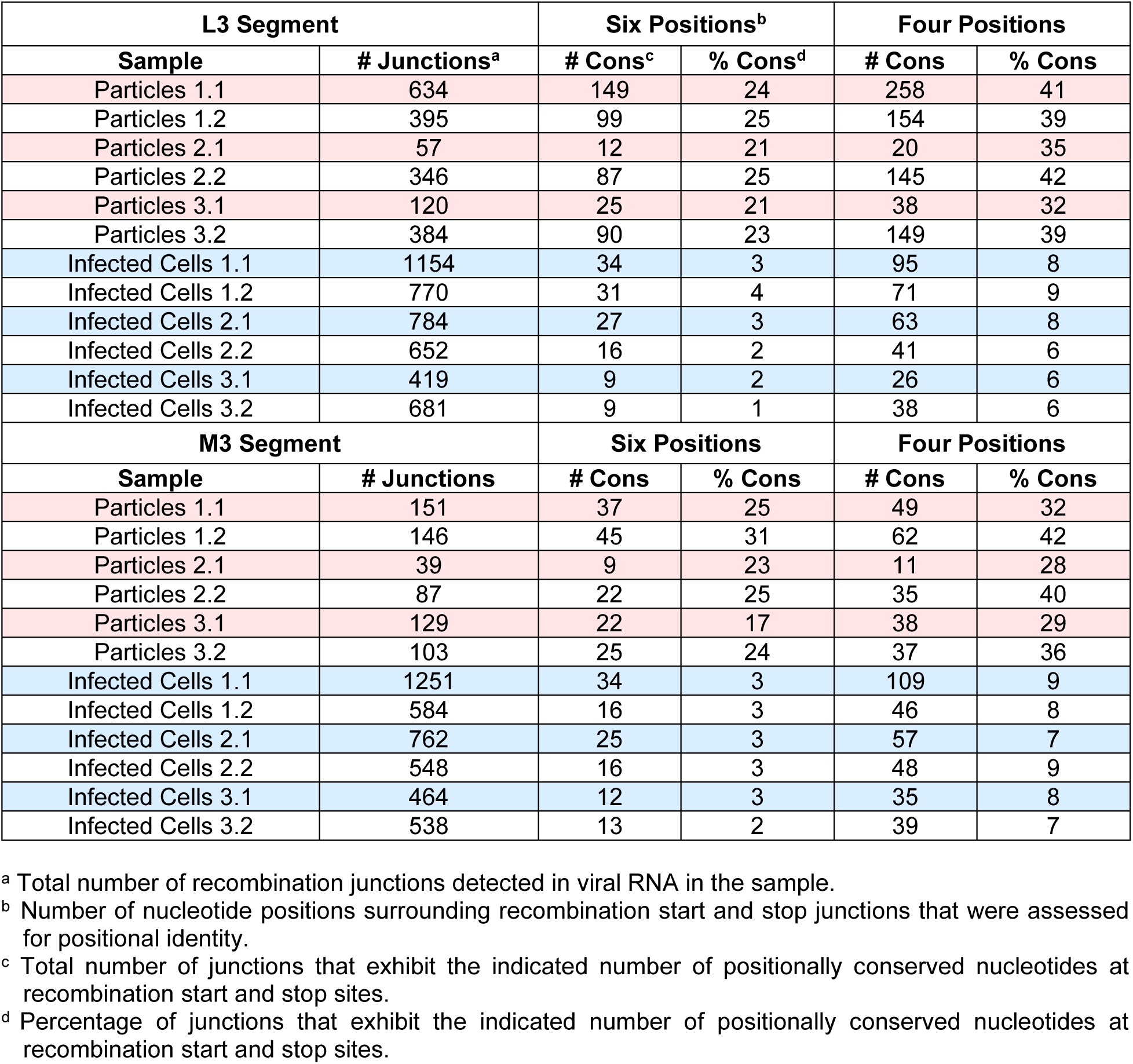
Microhomology surrounding junctions within the L3 and M3 segments.

Our findings are consistent with a model in which selection occurs during recombination and downstream processes, with the recombination landscape in virions representing a selectively filtered set of the population generated during infection. Differences in selection are reflected in the varied JFreqs between sample types (**S1 Table**), with the higher JFreq of intracellular viral RNA suggesting reduced recombination purification. Although potential recombination sites are similarly abundant, a small set of these sites is utilized in particle-associated RNA, whereas intracellular viral RNA exhibits more widespread use of available microhomology sequences (**Figs. 3 and S4**). Studies in other RNA virus systems have demonstrated differences between intracellular defective RNA populations and those incorporated into virions, implicating genome packaging as a key selective step [25, 28, 32]. For example, for influenza A virus, comparison of intracellular deletion viral genomes to those in released virions reveals a bottleneck during packaging; for alphaviruses, only a subset of defective RNAs generated in infected cells are packaged [25, 28]. Mechanisms of selective recombination and packaging remain to be determined.

## Materials and Methods

### Cell culture

L cells, obtained from Dr. Terence Dermody (University of Pittsburgh Medical Center), were propagated in suspension culture in Joklik’s minimum essential medium (JMEM; U.S. Biological) supplemented with 5% fetal bovine serum (FBS; Gibco), 2mM L-glutamine (Corning), and 100 U/ml penicillin plus 100 µg/ml streptomycin (Corning). Baby hamster kidney cells expressing T7 RNA polymerase under control of a cytomegalovirus promoter (BHK-T7), also provided by Dr. Terence Dermody, were maintained in Dulbecco’s minimum essential medium (DMEM; Corning) containing 10% tryptose-phosphate broth, 1% nonessential amino acids, 5% FBS, 2 mM L-glutamine, and 100 U/mL penicillin plus 100 µg/mL streptomycin. Geneticin (Gibco) was included at 1 mg/ml every other passage. All cells were incubated at 37°C in a humidified atmosphere containing 5% CO_2 a_nd were routinely screened for mycoplasma contamination.

### Viruses

Reovirus strain T1L was recovered using a plasmid-based reverse genetics system, supplied by Dr. Terence Dermody [33, 34]. Virus rescue was conducted in monolayers of BHK-T7 cells at approximately 50% confluency in 12-well plates. BHK-T7 cells were co-transfected with 0.8 µg of pT7 plasmids encoding each of the ten T1L reovirus positive-sense gene segments using TransIT-LT1 transfection reagent (Mirus Bio LLC) [33, 34]. A C3P3 plasmid, which was provided by Eukarÿs SAS, was utilized to increase efficiency [35]. Five days after transfection, virus was recovered by subjecting the cells to two cycles of freeze-thaw lysis. The recovered reovirus was plaque purified in L cells, and individual plaques were isolated and amplified twice in L cells to generate working virus stock [36].

### RNA collection and reovirus particle enrichment

4 × 10^8^ L cells were infected with three independent reovirus clones at a multiplicity of infection of 10 plaque forming units per cell (PFU/cell) at 37°C for 24 h. Following infection, a 10 ml aliquot of the infected cell suspension was collected for intracellular RNA analysis, while the remaining cells were processed for purification of viral particles. Total intracellular RNA was isolated using TRIzol Reagent (Invitrogen), according to the manufacturer’s instructions. Residual genomic DNA was removed by treatment with RNase-free DNase I (New England Biolabs) at 37°C for 10 minutes, followed by a second RNA purification using TRIzol LS Reagent (Invitrogen). Ribosomal RNA was subsequently depleted using the NEBNext rRNA Depletion Kit (New England Biolabs) following the manufacturer’s protocol.

Viral particles were purified from the remaining infected cells by deoxycholate permeabilization, Vertrel XF (DuPont) extraction, and CsCl gradient centrifugation, as described previously [17, 36]. Particle concentration was estimated using the conversion that one OD_260 u_nit corresponds to 2.1 × 10^12^ reovirus particles. For each replicate, 2.2 × 10^12^ viral particles were treated with benzonase (250 U) to remove extra-particle nucleic acids, and the reaction was terminated by the addition of 0.5 M EDTA (pH 8.0). Viral RNA was then extracted with Trizol LS Reagent (Invitrogen), according to the manufacturer’s protocol. The quality of all RNA samples was determined using a Bioanalyzer (Agilent), and the RNA concentration was determined by Qubit (Invitrogen).

### ClickSeq library preparation and next-generation sequencing (NGS)

Sequencing libraries were prepared from 250 ng of RNA per sample using the ClickSeq approach [29]. Briefly, reverse transcription was conducted with SuperScript III reverse transcriptase (Invitrogen) in the presence of 3′-azido-nucleotides at an AzNTP:dNTP ratio of 1:35, resulting in stochastic chain termination and generation of 3′-azido-blocked cDNA fragments. These cDNA products were subsequently ligated to Illumina i5 sequencing adaptors by click chemistry, followed by PCR amplification using barcoded Illumina p7 adaptors, [29]. Libraries were sequenced by 150-bp paired-end sequencing on a NovaSeq X Plus sequencing system (Illumina). Library quality control and next-generation sequencing were performed by ClickSeq Technologies LLC.

### Illumina RNA-sequencing processing and alignment

Sequencing reads were processed by trimming the Illumina TruSeq adapter sequences, reads shorter than 50 bp or containing bases with PHRED quality scores below 20 were removed using *fastp* [37]. The filtered FASTQ files were then mapped to the T1L reference genome (NCBI or GenBank accession numbers AF378003.1, AF129820.1, AF461682.1, AF490617.1, AF174382.1, M14779.1, L19774.1, M14325.1, M13139.1; the T1L L1 sequence differs from the GenBank reference and is available upon request) using the Python3 script *ViReMa* (Viral Recombination Mapper, version 0.32) [31] and the command line parameters python3 ViReMa.py reference_index input.fastq output.sam – Ouput_Dir sample_virema/--Output_Tag sample_virema-BED12 –MicroIndel_Length 5 –Defuzz 0. The sequence alignment map (SAM) files generated were subsequently processed with the samtools [38] suite to calculate nucleotide depth at each position in a sorted binary alignment map (BAM) file using command line parameters samtools depth -a -m 0 sample_virema.sorted.bam > sample_virema.coverage.

### Recombination junction analysis

ViReMa output files were filtered to separate recombination junctions containing microinsertions or microdeletions of ≤5 nt into independent BED files. The frequency of the start position of each junction was evaluated by dividing the frequency of reads mapping to a recombination junction by the total mapped reads as previously described [10].

### Principal coordinate analysis

A unique ID was created for each recombination event using the *dplyr* package in RStudio [39]. All unique recombination events were identified for each virus, and the frequency value was added to a matrix. NA values were substituted with zero. Bray-Curtis dissimilarity matrix was generated using the vegdist function in the *vegan* package. PCoA was employed to ordain the samples. Data were visualized using *ggplot2*, with the *ggrepel* package used to prevent label overlap.

### Nucleotide composition analysis and sequence microhomology quantification

Nucleotide composition surrounding recombination junctions was analyzed at both junction start and stop sites. To minimize bias introduced by highly abundant recombination events, each unique junction was weighted equally, regardless of read count [32]. Sequences were extracted from sorted BED files, converted to a text format, and processed in RStudio. Start site and stop site sequences were separated and stop site sequences containing less than 51 nucleotides were excluded from further analysis. Position-specific nucleotide frequencies were calculated using the *Biostrings* package [40]. The nucleotide frequency was plotted as a logoplot using the *ggseqlogo* package in RStudio [41]. For microhomology quantification, the nucleotides surrounding the junction sites were compared for positional identity relative to the start and stop sites. Positionally identical sequences were quantified in RStudio and plotted in GraphPad Prism 10 as previously described [10].

### Microhomology sequence occurrence and usage

Microhomology frequencies were quantified by extracting symmetric nucleotide windows (four or six nucleotides) surrounding each recombination junction using the *dplyr* package [39]. Corresponding sequences from the junction start and stop sites were compared to determine the extent of positional nucleotide identity, and the percentage of recombination junctions containing microhomology of each length was calculated using the *Biostrings* package in RStudio.

To evaluate the genomic distribution of conserved four-nucleotide microhomology, unique four-nucleotide sequences between the start and stop sites of recombination junctions within the L1, L3, M1, and M3 gene segments were extracted using *dplyr* [39]. Reverse-complement sequences were generated for junctions originating from the minus strand using *Biostrings* [40]. Each conserved four-nucleotide sequence was then mapped back to the corresponding T1L reference gene segment, and all genomic occurrences were identified using *Biostrings* [40]. The location of each short sequence in the viral genome was extracted using *Biostrings* [40]. Genomic coordinates were adjusted to correspond with the recombination start and stop reported by *ViReMa*, and the number of occurrences associated with each junction start and stop site was quantified for downstream analyses.

### Statistical analyses

All statistical analyses were performed using GraphPad Prism version 10. The statistical tests applied to each dataset are specified in the corresponding figure legends. Selection of the appropriate statistical tests was made in consultation with a biostatistician.

## Data Availability Statement

Data generated by ClickSeq sequencing can be accessed at NCBI Sequence Read Archive (SRA) under the BioProject accession PRJNA1519998. Code utilized in this report can be accessed at (https://github.com/ogdenlab1/ReoRecombination).

## Acknowledgements

We thank Dr. James C. Slaughter for biostatistics expertise.

