## Supplementary Tables and Figures for "Comparison of intracellular and encapsidated viral RNA reveals selective constraints on reovirus recombination"

#### S1 Table. Alignment statistics

| Alignment statistic | Segment | Particles 1.1 | Particles 2.1 | Particles 3.1 | Infected Cells 1.1 | Infected Cells 2.1 | Infected Cells 3.1 |
|---|---|---|---|---|---|---|---|
| **Experiment 1** | | | | | | | |
| **Total reads** | | 106,937,620 | 201,517,204 | 160,372,824 | 109,422,168 | 11,610,090 | 129,575,404 |
| **Total reads mapped to virus** | | 84,652,206 | 121,649,536 | 145,511,412 | 81,993,948 | 6,859,099 | 24,082,593 |
| **% reads mapped to virus** | | 77.4% | 60.4% | 90.7% | 74.9% | 59.1% | 18.6% |
| **Jfreq [a]** | | 6.05 | 10.8 | 11.7 | 42.9 | 43.1 | 52.1 |
| **# reads with recombination events** | | | | | | | |
| | L1 | 12,503 | 32,665 | 32,483 | 23,491 | 2,193 | 10,053 |
| | L2 | 9,013 | 16,960 | 14,760 | 24,328 | 1,875 | 9,207 |
| | L3 | 11,209 | 10,130 | 16,859 | 41,883 | 4,511 | 18,074 |
| | M1 | 1,636 | 7,454 | 16,605 | 20,338 | 1,574 | 7,707 |
| | M2 | 4,754 | 3,621 | 11,692 | 49,834 | 4,064 | 18,943 |
| | M3 | 2,764 | 18,586 | 49,702 | 49,423 | 4,461 | 19,966 |
| | S1 | 4,420 | 24,818 | 17,309 | 42,315 | 3,250 | 14,343 |
| | S2 | 1,375 | 2,538 | 4,050 | 29,317 | 2,269 | 9,048 |
| | S3 | 672 | 12,046 | 4,992 | 38,228 | 2,833 | 9,967 |
| | S4 | 1,711 | 2,837 | 2,315 | 32,282 | 2,514 | 8,076 |
| **% plus-strand recombination event reads** | | 39 | 52 | 54 | 99 | 99 | 99 |
| **# reads mapped to viral gene segment** | | | | | | | |
| | L1 | 17,174,296 | 20,849,740 | 22,714,366 | 5,160,999 | 612,022 | 2,045,068 |
| | L2 | 15,931,333 | 16,988,248 | 14,542,591 | 5,384,298 | 490,706 | 1,726,465 |
| | L3 | 17,156,344 | 24,993,910 | 22,460,002 | 7,278,225 | 892,113 | 3,064,824 |
| | M1 | 5,894,789 | 15,812,151 | 27,942,353 | 5,718,800 | 482,358 | 1,486,982 |
| | M2 | 9,845,200 | 6,686,857 | 8,191,612 | 8,094,865 | 726,353 | 2,303,701 |
| | M3 | 4,797,993 | 13,329,124 | 19,930,153 | 13,049,918 | 921,850 | 3,851,429 |
| | S1 | 4,962,942 | 9,651,183 | 11,985,374 | 12,037,142 | 906,270 | 3,407,273 |
| | S2 | 3,518,887 | 5,338,459 | 5,519,932 | 7,264,720 | 545,484 | 1,783,803 |
| | S3 | 2,210,439 | 5,243,670 | 7,891,616 | 7,872,710 | 525,665 | 1,748,003 |
| | S4 | 3,159,983 | 2,756,194 | 4,333,413 | 10,132,271 | 756,278 | 2,665,045 |
| **Experiment 2** | | | | | | | |
| **Alignment statistic** | **Segment** | **Particles 1.2** | **Particles 2.2** | **Particles 3.2** | **Infected Cells 1.2** | **Infected Cells 2.2** | **Infected Cells 3.2** |
| **Total reads** | | 60,888,632 | 64,320,894 | 105,147,916 | 66,510,566 | 56,447,898 | 68,424,616 |
| **Total reads mapped to virus** | | 59,640,790 | 59,152,634 | 99,461,794 | 46,006,167 | 35,070,858 | 41,421,736 |
| **% reads mapped to virus** | | 98.0% | 92.0% | 94.6% | 69.2% | 62.1% | 62.5% |
| **Jfreq [a]** | | 4.53 | 15.11 | 7.47 | 35.39 | 49.03 | 54.04 |
| **# reads with recombination events** | | | | | | | |
| | L1 | 6,605 | 15,022 | 17,067 | 11,576 | 11,845 | 19,201 |
| | L2 | 2,970 | 18,793 | 9,261 | 9,682 | 13,904 | 19,998 |
| | L3 | 5,832 | 32,842 | 17,056 | 29,285 | 26,468 | 41,734 |
| | M1 | 1,999 | 6,238 | 4,558 | 8,985 | 10,341 | 15,188 |
| | M2 | 1,026 | 672 | 8,828 | 25,535 | 27,125 | 30,650 |
| | M3 | 1,578 | 8,152 | 4,613 | 21,683 | 23,271 | 33,671 |
| | S1 | 4,864 | 4,360 | 6,013 | 17,033 | 17,128 | 20,642 |
| | S2 | 405 | 784 | 2,043 | 10,369 | 11,325 | 16,869 |
| | S3 | 1,294 | 1,314 | 1,574 | 13,629 | 15,179 | 15,146 |
| | S4 | 448 | 1,204 | 3,323 | 15,047 | 15,357 | 17,904 |
| **% plus-strand recombination event reads** | | 41 | 47 | 36 | 98 | 98 | 98 |
| **# reads mapped to viral gene segment** | | | | | | | |
| | L1 | 9,542,430 | 10,743,996 | 17,372,463 | 3,932,584 | 3,185,489 | 3,727,342 |
| | L2 | 8,655,838 | 9,797,740 | 14,772,893 | 3,467,987 | 2,670,714 | 3,469,121 |
| | L3 | 10,716,236 | 16,086,912 | 19,292,544 | 6,966,150 | 4,791,808 | 5,735,751 |
| | M1 | 4,213,219 | 5,353,444 | 8,980,184 | 3,220,862 | 2,445,488 | **3,026,597** |
| | M2 | 4,163,832 | 1,956,934 | 13,835,632 | 5,040,323 | 3,752,893 | 4,214,610 |
| | M3 | 4,861,078 | 4,326,413 | 5,641,157 | 5,956,914 | 4,087,881 | 5,929,270 |
| | S1 | 9,566,557 | 5,397,665 | 6,844,393 | 5,555,558 | 4,676,863 | 5,108,730 |
| | S2 | 1,855,411 | 1,656,766 | 4,974,141 | 3,007,232 | 2,466,683 | 3,111,154 |
| | S3 | 4,901,957 | 2,689,384 | 2,855,734 | 3,066,173 | 2,160,259 | 2,554,189 |
| | S4 | 1,164,232 | 1,143,380 | 4,892,653 | 5,792,384 | 4,832,780 | 4,544,972 |

[a] Number of reads containing junctions per 10,000 mapped reads

**S2 Table. Microhomology surrounding junctions within the L1 and M1 segments.**

| L1 Segment | | Six Positions | | Four Positions | |
|---|---|---|---|---|---|
| Sample | # Junctions | # Cons | % Cons | # Cons | % Cons |
| Particles 1.1 | 716 | 171 | 24 | 310 | 43 |
| Particles 1.2 | 370 | 104 | 28 | 164 | 44 |
| Particles 2.1 | 55 | 11 | 20 | 21 | 38 |
| Particles 2.2 | 166 | 54 | 33 | 76 | 46 |
| Particles 3.1 | 100 | 21 | 21 | 36 | 36 |
| Particles 3.2 | 351 | 86 | 25 | 146 | 42 |
| Infected Cells 1.1 | 558 | 18 | 3 | 36 | 6 |
| Infected Cells 1.2 | 280 | 14 | 5 | 32 | 11 |
| Infected Cells 2.1 | 377 | 8 | 2 | 28 | 7 |
| Infected Cells 2.2 | 269 | 9 | 3 | 29 | 11 |
| Infected Cells 3.1 | 209 | 6 | 3 | 19 | 9 |
| Infected Cells 3.2 | 316 | 12 | 4 | 25 | 8 |
| **M1 Segment** | | **Six Positions** | | **Four Positions** | |
| **Sample** | **# Junctions** | **# Cons** | **% Cons** | **# Cons** | **% Cons** |
| Particles 1.1 | 121 | 32 | 26 | 46 | 38 |
| Particles 1.2 | 133 | 38 | 29 | 62 | 47 |
| Particles 2.1 | 24 | 4 | 17 | 7 | 29 |
| Particles 2.2 | 73 | 21 | 29 | 29 | 40 |
| Particles 3.1 | 107 | 32 | 30 | 47 | 44 |
| Particles 3.2 | 120 | 38 | 32 | 49 | 41 |
| Infected Cells 1.1 | 585 | 14 | 2 | 49 | 8 |
| Infected Cells 1.2 | 272 | 9 | 3 | 19 | 7 |
| Infected Cells 2.1 | 314 | 5 | 2 | 31 | 10 |
| Infected Cells 2.2 | 271 | 5 | 2 | 21 | 8 |
| Infected Cells 3.1 | 158 | 4 | 3 | 11 | 7 |
| Infected Cells 3.2 | 259 | 6 | 2 | 20 | 8 |

[a] Total number of recombination junctions detected in viral RNA in the sample.

[b] Number of nucleotide positions surrounding recombination start and stop junctions that were assessed for positional identity.

[c] Total number of junctions that exhibit the indicated number of positionally conserved nucleotides at recombination start and stop sites.

[d] Percentage of junctions that exhibit the indicated number of positionally conserved nucleotides at recombination start and stop sites.

### SUPPLEMENTARY FIGURES

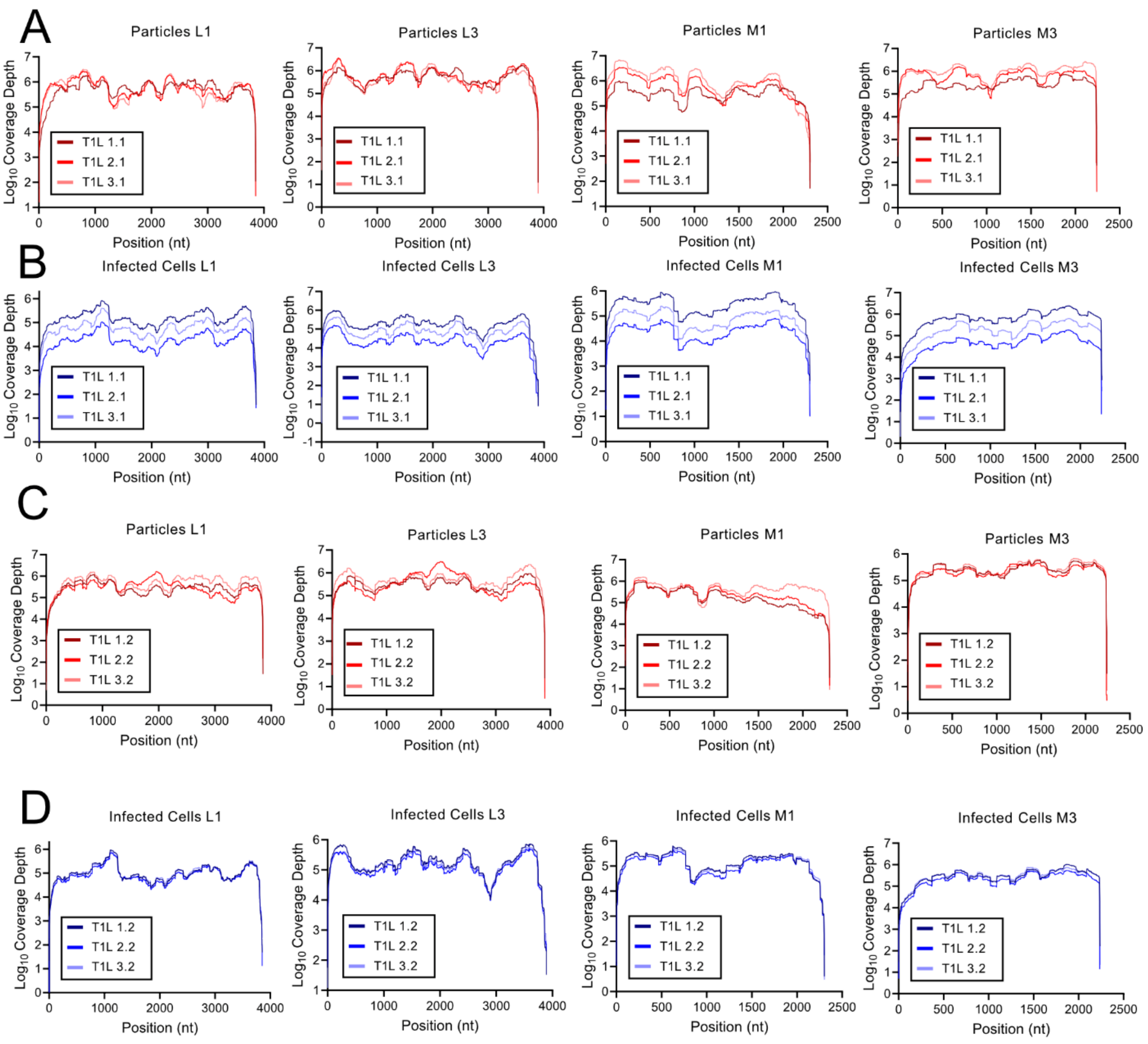


**S1 Fig. Reovirus sequence coverage across genome segments.** Sequence read depth across all nucleotide positions for segments L1, L3, M1, and M3 from experiment 1 (A-B) and experiment 2 (C-D). Coverage for three individual virus clones each in particle samples (A, C) and infected-cell samples (B, D) is shown.

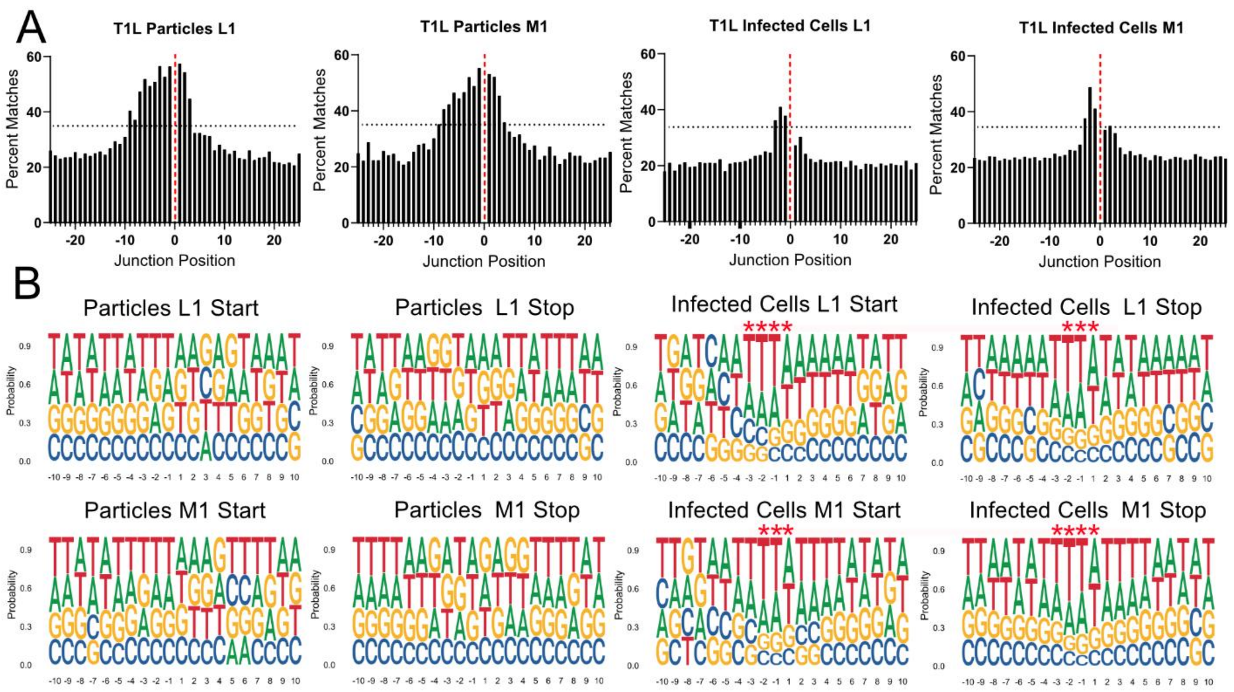


**S2 Fig. Microhomology and logoplots.** (A) Sequence microhomology surrounding recombination junctions in particle and infected-cell viral RNA. The nucleotides that overlap at each position in 50-bp regions flanking the start and stop recombination junction sites were quantified for segments L1 and M1 and are expressed as a percentage of the total detected recombination junctions. The junction is labeled zero and denoted with a dashed red line. The black dashed line denotes y = 35. (B) Logoplots showing nucleotides surrounding recombination junctions in particle and infected-cell viral RNA. Nucleotide composition was calculated as the percent adenosine (A), cytosine (C), guanine (G), and thymine (T) at each position in a 20-bp region surrounding the start and stop recombination junction sites. Positions upstream (−10 to −1) and downstream (+1 to +10) of the junction position are indicated. Red asterisks denote statistical significance via a Chi-Square Goodness-of-Fit test against an expected uniform baseline distribution and a threshold at which the dominant nucleotide constitutes at least 35% of the total local population. Each analyzed condition contains combined data from two independent experiments, each comprising three biological replicates. Sequence logo and microhomology analyses were performed using all unique recombination junctions detected in each condition: L1 particles (n = 1,758), L1 infected cells (n = 2,009), M1 particles (n = 578), and M1 infected cells (n = 1,859).

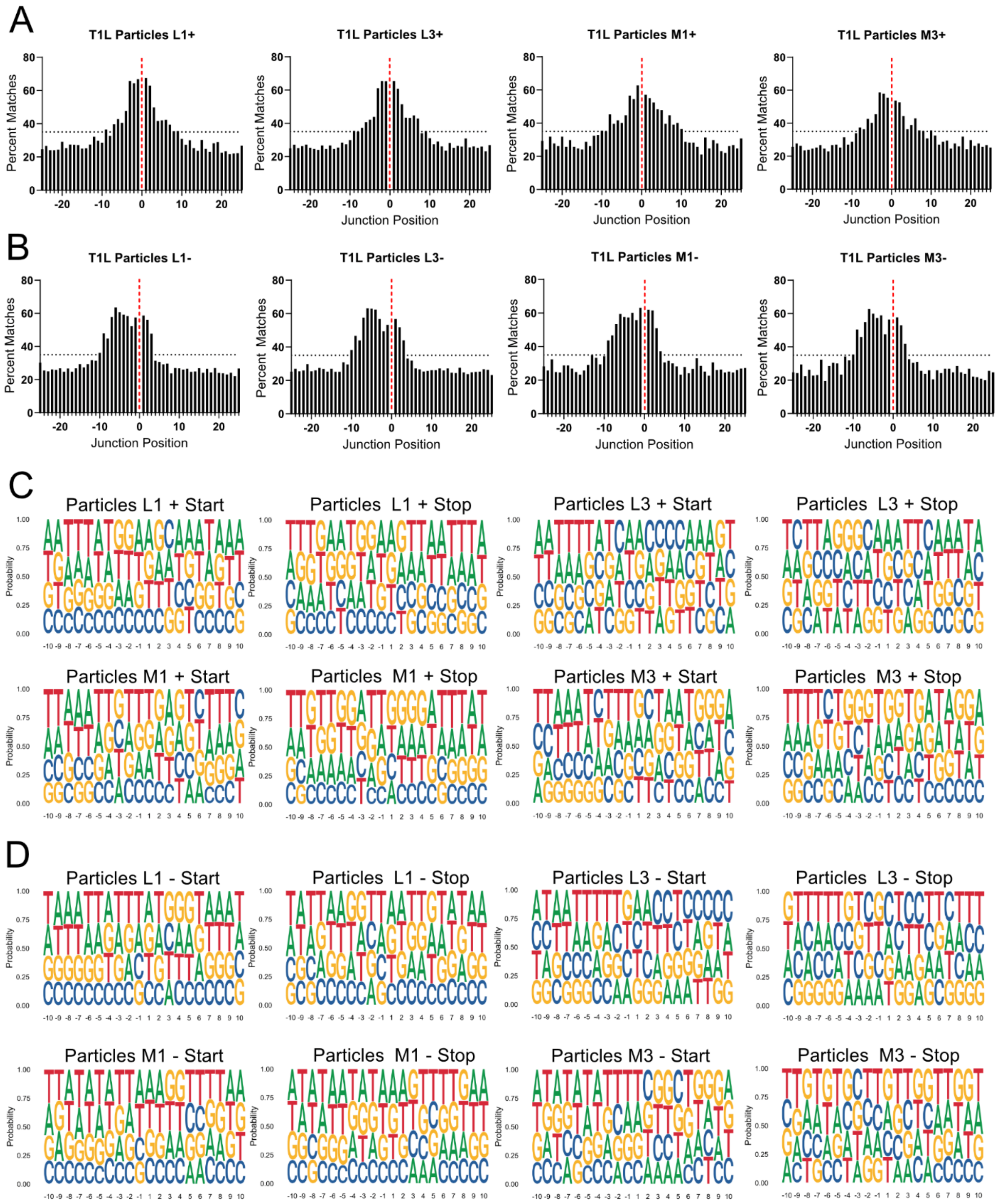


**S3 Fig. Sequence microhomology and logoplots surrounding recombination junctions in particle-encapsidated viral RNA by strand orientation.** (A-B) Microhomology surrounding recombination junctions in particle-encapsidated viral RNA was analyzed separately for the plus strand (A) and minus strand (B). The nucleotides overlapping at each position within 50-bp regions flanking the start and stop recombination junction sites were quantified for the L1, L3, M1, and M3 gene segments and are expressed as the percentage of total detected recombination junctions. The recombination junction is designated as position zero and indicated by the dashed red line. The black dashed line denotes y = 35. (C-D) Logoplots showing nucleotides surrounding recombination junctions in the plus strand (C) and minus strand (D). Nucleotide composition was calculated as the percent adenosine (A), cytosine (C), guanine (G), and thymine (T) at each position in a 20-bp region surrounding the start and stop recombination junction sites. Positions upstream (−10 to −1) and downstream (+1 to +10) of the junction position are indicated. Red asterisks denote statistical significance calculated using a Chi-Square Goodness-of-Fit test against an expected uniform baseline distribution and a threshold at which the dominant nucleotide constitutes at least 35% of the total local population. Each analysis includes combined data from two independent experiments, each consisting of three biological replicates. Microhomology and sequence logo analyses were conducted using all unique recombination junctions detected for each strand: L1 plus strand (n = 613), L3 plus strand (n = 908), M1 plus strand (n = 194), M3 plus strand (n = 288), L1 minus strand (n = 1,047), L3 minus strand (n = 911), M1 minus strand (n = 329), and M3 minus strand (n = 315).

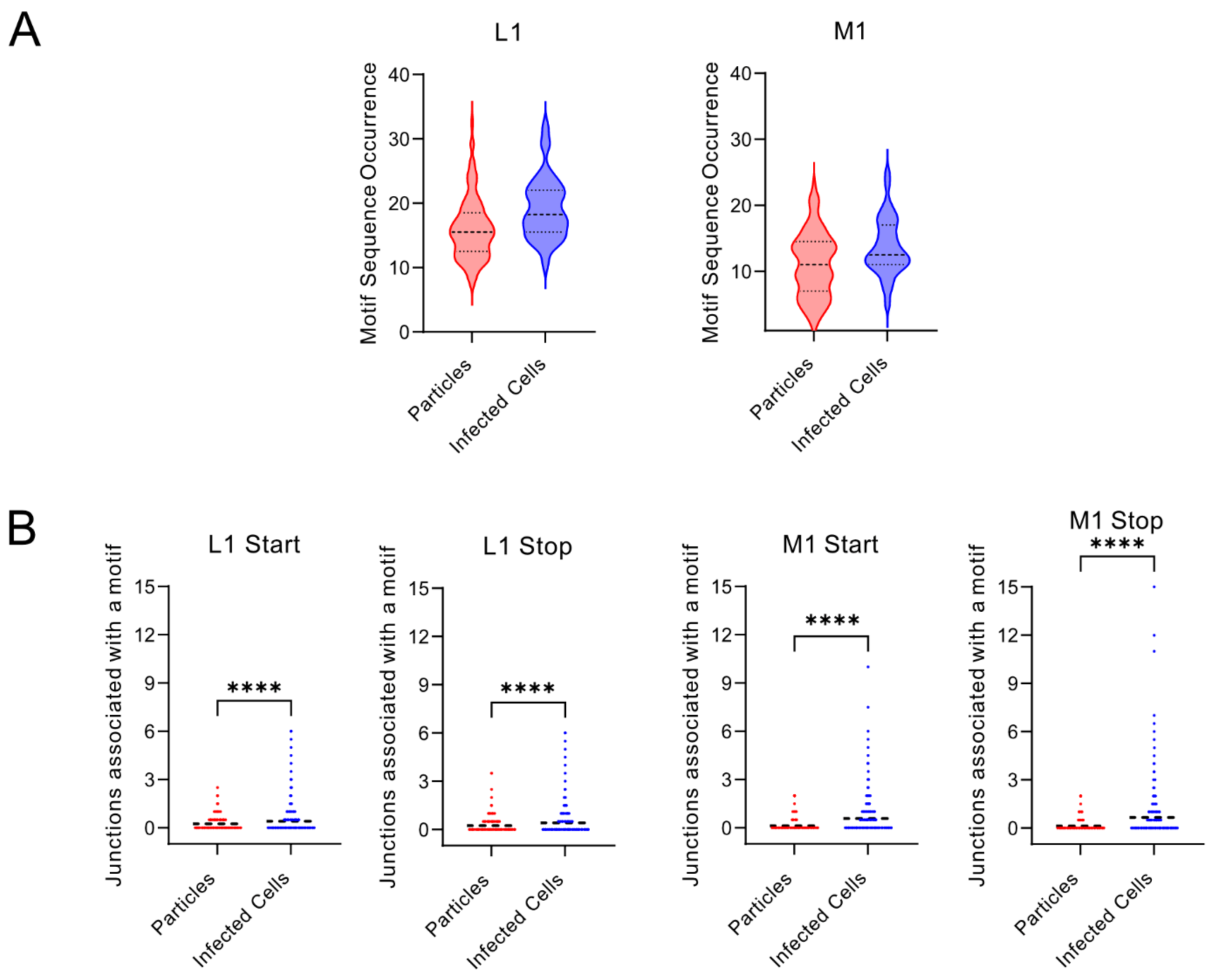


**S4 Fig. Recombination site selection.** (A) Frequency of microhomology sequences in segments. The number of times each unique four-nucleotide microhomology sequence identified at a recombination junction is present in segment L1 or M1 was computationally quantified. For particles, L1: n = 355 and M1: n = 164 unique four-nucleotide microhomology sequences. For infected cells, L1: n = 115 and M1: n = 98 unique four-nucleotide microhomology sequences. Dashed lines represent the median and quartiles. (B) Usage of microhomology sites for recombination. The number of times each occurrence of a four-nucleotide sequence from a unique start site (B) or stop site (C) analyzed in (A) was incorporated into a recombination site in the L1 or M1 segment was computationally quantified for particles and infected cells. Each analyzed condition contains combined data from two independent experiments, each comprising three biological replicates. For particles, L1: n= 5670 and M1: n= 1826 instances. For infected cells, L1: n= 2198 and M1: n= 1311 instances. Dashed lines represent the mean.
